# Towards Analyzing the Reclassification Dynamics of ClinVar Variants

**DOI:** 10.1101/2023.01.24.525342

**Authors:** Andreas Ruscheinski, Anna Lena Reimler, Roland Ewald, Adelinde M. Uhrmacher

## Abstract

ClinVar aggregates information about genetic variants and their relation to human diseases and forms a valuable resource for clinical diagnostics. The assessment of variants, e.g., as being benign or pathogenetic, changes over time. We collected variant classification histories of ClinVar releases and used those to derive discrete-time Markov chains to investigate the reclassification dynamics of variants in ClinVar in terms of transition probabilities between ClinVar releases and reclassifications.

## 1 Introduction

ClinVar is the most comprehensive public data source for the clinical interpretation of genetic variants in the human genome [4]. ClinVar is kept up to date by geneticists, who continuously submit variant interpretations backed by real-world clinical evidence. A variant interpretation classifies a genetic variant according to the ACMG guidelines [5] with a significance of either *pathogenic* (`p`) (i.e., disease-causing), *likely pathogenic* (`lp`), *uncertain* (`u`), *likely benign* (`lb`), or *benign* (`b`). The submitted variant interpretations are then aggregated, curated, and complemented with supporting evidence by researchers, clinical laboratories, and expert panels [4]. Suppose new submissions offer clinical evidence that contradicts the current interpretation of a variant. In that case, the variant gets *reclassified*. Since ClinVar submissions may disagree with each other on the interpretation of a variant, ClinVar introduces additional ‘aggregate significances’ to distinguish different levels of disagreement. In case of a slight disagreement, i.e., `b` vs. `lb` or `p` vs. `lp`, the variant gets classified as *benign/likely benign* or *pathogenic/likely pathogenic* whereas strong disagreements (e.g., `b` vs. `p`) are classified as *conflicting interpretations of pathogenicity* (`c`).

There are several reasons why ClinVar variant reclassifications are an interesting phenomenon to study. Firstly, a reclassification may motivate clinicians to reevaluate medical cases where the variant has been found, i.e., to check whether this change needs to be reported to the patient (e.g. in the form of an updated diagnostic report). Understanding the reclassification dynamics thus allows us to estimate the additional effort clinicians and genetic labs have to reserve for this work. Secondly, ClinVar is often used as a benchmark (‘ground truth’) to evaluate AI-based methods for variant interpretation [3]. Understanding both the frequency and overall impact of reclassifications— and any biases this may introduce — also allows us to better understand the limitations of such benchmarking against imperfect data. Thirdly, the ClinVar project is a large-scale, global endeavour to compile clinical genetics knowledge over a long time horizon. Analyzing the current dynamics of variant reclassification may thus help to pinpoint potential areas of improvement and spot weaknesses in the current curation process.

Previous work [6,2] compared the classification of variants between selected ClinVar releases focusing on variants classified as likely benign and likely pathogenic to investigate the uncertainty in variant interpretation. In contrast, our approach takes into account the entire classification histories of all Clinvar variants to derive a discrete-time Markov chain capturing the reclassification dynamics.

## 2 From ClinVar variants to a Markov Chain

Our key idea is to treat classification histories of the variants as possible sequences generated by an unknown stochastic process that, once known, can be analyzed to gain insights into reclassification dynamics. Here, we assume that the stochastic process can be modelled as a Markov chain according to which the probability of the subsequent classification only depends on the present classification of the variant [1]. To describe such a Markov chain, we have to estimate the transition probabilities between the different classifications using the histories of the variants. To simplify our analysis, we will not take into account slight disagreements but simply treat them as `b` or `p` respectively, which leaves us with `b`, `p`, `u` and `c` as categories respectively.

To this end, we aggregated the classification histories of all variants from all ClinVar releases and represented them as strings, where each character represents a classification of a variant in a specific ClinVar release. Next, we ex-cluded the variants which were not reclassified.^3^ Afterwards, we counted the pair-wise adjacent occurrences of classifications in the strings. Finally, we estimated the transition probabilities using the maximum likelihood estimator 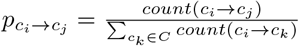, where *c_i_, C_j_* ∈ ***C*** and ***C*** is the set of possible classifications of a variant. The approach is illustrated based on a single variant history in Figure 1.

**Fig. 1.**
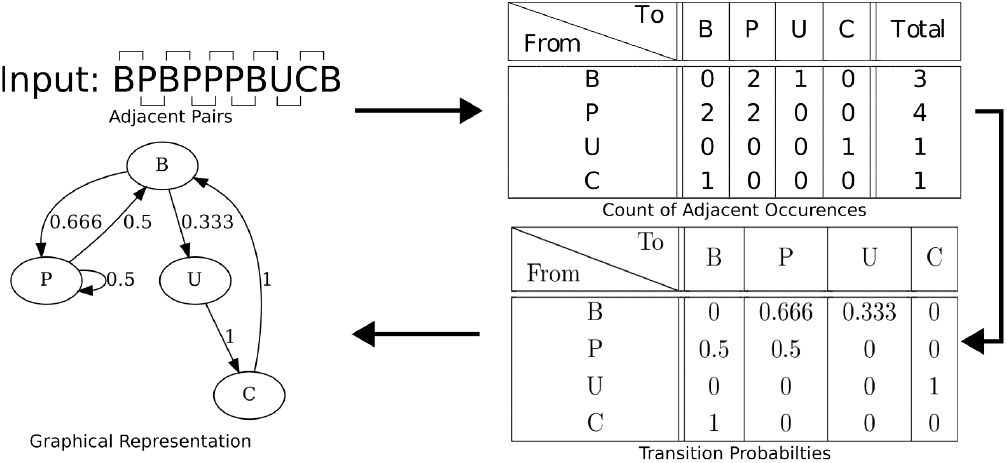
Deriving a Markov chain from a variant’s history

We implemented a data pipeline that automatically downloads all ClinVar releases and encodes the variant classifications into strings, with each character representing the classification of a variant in a given ClinVar release. In total, we processed 1, 496, 916 variants of which 91, 830 (≈6%) were reclassified. Altogether we considered 17, 094, 260 classifications with 110, 802 reclassification events. Counting these events allows us to estimate the transition probabilities of the Markov chain.

Considering the distribution of reclassifications per variant in figure 2, we see that most reclassified variants have been reclassified just once. The figure also shows that the number of variants with at least ***n*** reclassification events is exponentially distributed (note the log scale on the y axis).

**Fig. 2.**
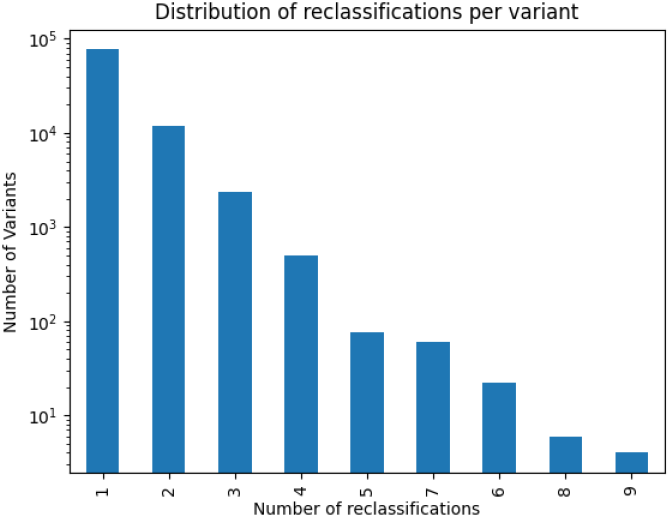
Distribution of reclassifications per variant

Figure 3 shows the Markov chain with all transition probabilities. Reclassifications are relatively unlikely, i.e. most variants are not reclassified at all. The highest transition probability is from uncertain to conflicting (0.8%). Generally, transitions to the class *conflicting*, if not remaining in the old class, are the most likely ones, (transition probabilities of 0.7%, 0.8%, and 0.6% for b, u, and p respectively). In comparison, transitioning from *conflicting* to another class is less likely (0.1%). This indicates a need for potential future adjustments to the ClinVar review process: with the current dynamics, the number of variants with conflicting interpretations will otherwise continue to increase.

**Fig. 3.**
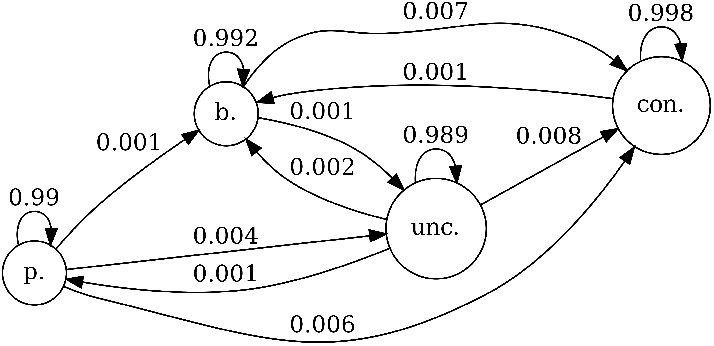
Markov-Chain showing the transition probabilities of classification between ClinVar releases (probabilities less then 10^-3^ ommitted)

Figure 4 shows the same data but only considers transitions between *different* states, i.e. re-classifications. We see that most *conflicting* variants are reclassified into the significance groups *benign* or *uncertain*; the probability that they will be classified as *pathogenic* is only 15%. Such insights may help practitioners in the later diagnosis.

**Fig. 4.**
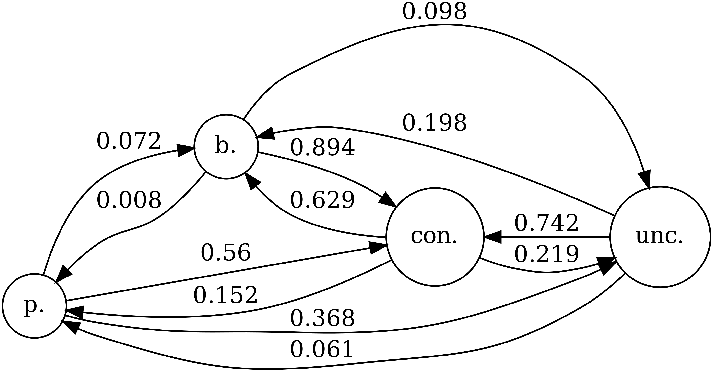
Markov-Chain showing the transition probabilities of reclassifications

## 3 Future work and Conclusion

Our analysis shows that the ClinVar curation approach generally works very well. Only a few variants (6%) ever get reclassified, and most reclassified variants only get reclassified *once*. However, we also found that the ClinVar curators might face a growing set of variants with conflicting interpretations if the current reclassification dynamics persist. ClinVar’s recent addition of new, non-ACMG significance types, such as *pathogenic, low-penetrance*, might be a reaction (and remedy) to this issue.

In the future, we plan to extend our analysis to see potential *changes* in the reclassification dynamics over time, i.e., to interpret variant histories as (discrete) time series. We also want to consider the differences in the transition probabilities when restricting the analysis to specific variants, e.g. those already reviewed by expert panels, those with particularly many submissions, or those that are challenging to detect with current sequencing technology.

Another avenue of future research could be to find common characteristics of particularly challenging variants. Such variants could also be a fruitful subset of ClinVar to benchmark AI-based approaches for variant interpretation.

## Supporting information

Poster

## Footnotes

3 Therefore, our Markov Chain only models the transition probabilities of variants that have been reclassified at least once, i.e., the 91.830 see below.

## References

1. Ames, C.: The markov process as a compositional model: A survey and tutorial. Leonardo 22(2), 175–187 (1989)

2. Harrison, S.M., Rehm, H.L.: Is ‘likely pathogenic’really 90% likely? reclassification data in clinvar. Genome medicine 11(1), 1–4 (2019)

3. Ioannidis, N.M., et al.: Revel: an ensemble method for predicting the pathogenicity of rare missense variants. The American Journal of Human Genetics 99(4), 877–885 (2016)

4. Landrum, M.J., Kattman, B.L.: Clinvar at five years: Delivering on the promise. Human mutation 39(11), 1623–1630 (2018)

5. Richards, S., et al.: Standards and guidelines for the interpretation of sequence variants: a joint consensus recommendation of the american college of medical genetics and genomics and the association for molecular pathology. Genetics in medicine 17(5), 405–423 (2015)

6. Xiang, J., et al.: Reinterpretation of common pathogenic variants in clinvar revealed a high proportion of downgrades. Scientific reports 10(1), 1–5 (2020)

