## Supplementary material for "Towards Analyzing the Reclassification Dynamics of ClinVar Variants": Poster

Traditio et Innovatio

### Motivation

- ClinVar collects and aggregates variant interpretations submitted by researchers, clinical laboratories, and expert panels
- ClinVar is the most comprehensive public data source for clinical interpretations of genetic variants in the human genome
- Clinical interpretations classify variants as pathogenic (p), likely pathogenic (lp), uncertain (u), likely benign (lb), or benign (b)
- When submissions contradict the current variant interpretation the variant get reclassified
    - Slight disagreement (p vs. lp or b vs. lb): pathogenic/likely pathogenic or benign/likely benign
    - Strong disagreement (p vs. b): conflicting interpretations of pathogenicity (c)

Reasons to study reclassification dynamics:

1. **Clinical Practice:** Estimate the additional effort clinicians and clinical labs have to reserve for reconsidering previous cases
2. **Machine Learning:** Understanding the limitations of using ClinVar for benchmarking AI-methods for variant interpretation
3. **Curation Process:** Pinpoint potential areas of improvement and spot weaknesses in current curation process

**Our approach:** Analyzing the reclassification dynamics via a discrete-time Markov chain build from the classification histories of all ClinVar variants

### From ClinVar variants to a Markov Chain

- **Assumption:** Next classification only depends on current classification → Markov chain
- **Method:**
    1. Build strings for all variants where the characters represent the classifications
    2. Count pair-wise adjacent occurrences of classifications in all strings
    3. Calculate transition probabilities using maximum-likelihood estimation

$$p_{c_i \to c_j} = \frac{count(c_i \to c_j)}{\sum_{c_k \in C} count(c_i \to c_k)}$$

Input: BPBPPPBUCB

Adjacent Pairs

| From \ To | B | P | U | C | Total |
|---|---|---|---|---|---|
| B | 0 | 2 | 1 | 0 | 3 |
| P | 2 | 2 | 0 | 0 | 4 |
| U | 0 | 0 | 0 | 1 | 1 |
| C | 1 | 0 | 0 | 0 | 1 |

Count of Adjacent Occurences

| From \ To | B | P | U | C |
|---|---|---|---|---|
| B | 0 | 0.666 | 0.333 | 0 |
| P | 0.5 | 0.5 | 0 | 0 |
| U | 0 | 0 | 0 | 1 |
| C | 1 | 0 | 0 | 0 |

Transition Probabilties

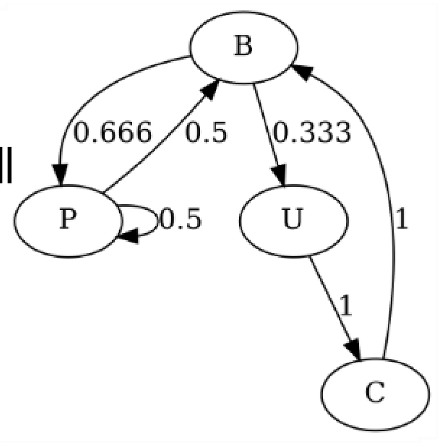


Graphical Representation

### Results

In total processed 1,496,916 variants of which 91,830 (6%) were reclassified

Fig. 1: Distribution of reclassifications per variant

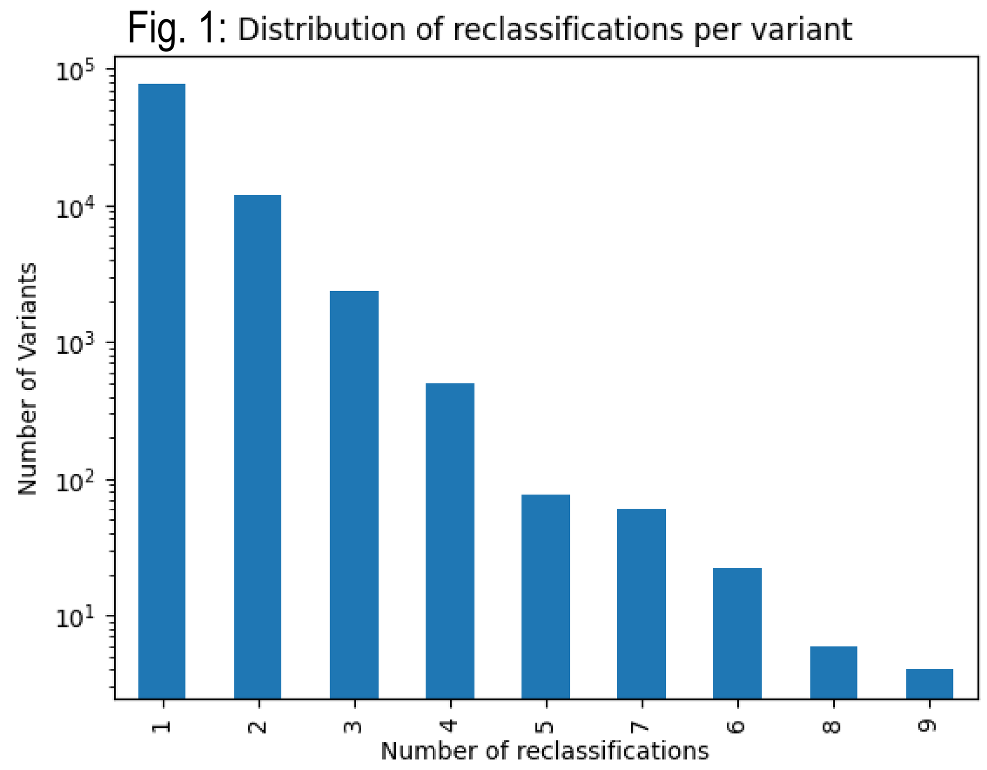


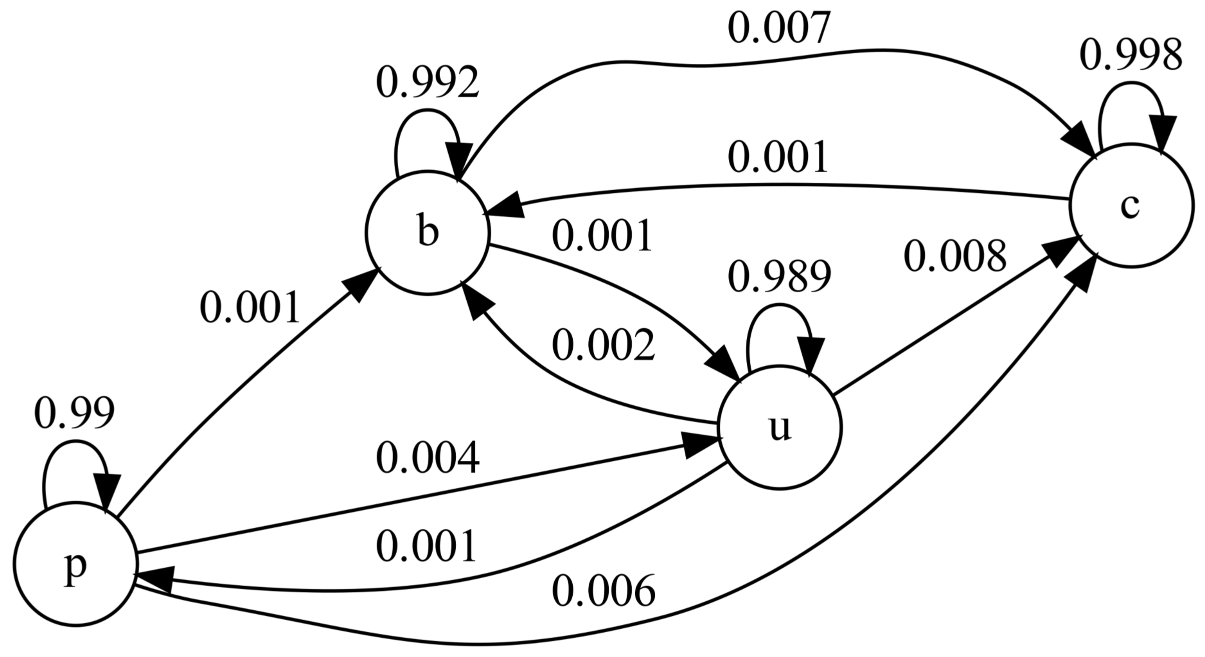


Fig. 2: Markov chain - Transition probabilities between ClinVar releases (probabilities less then 0.001 omitted)

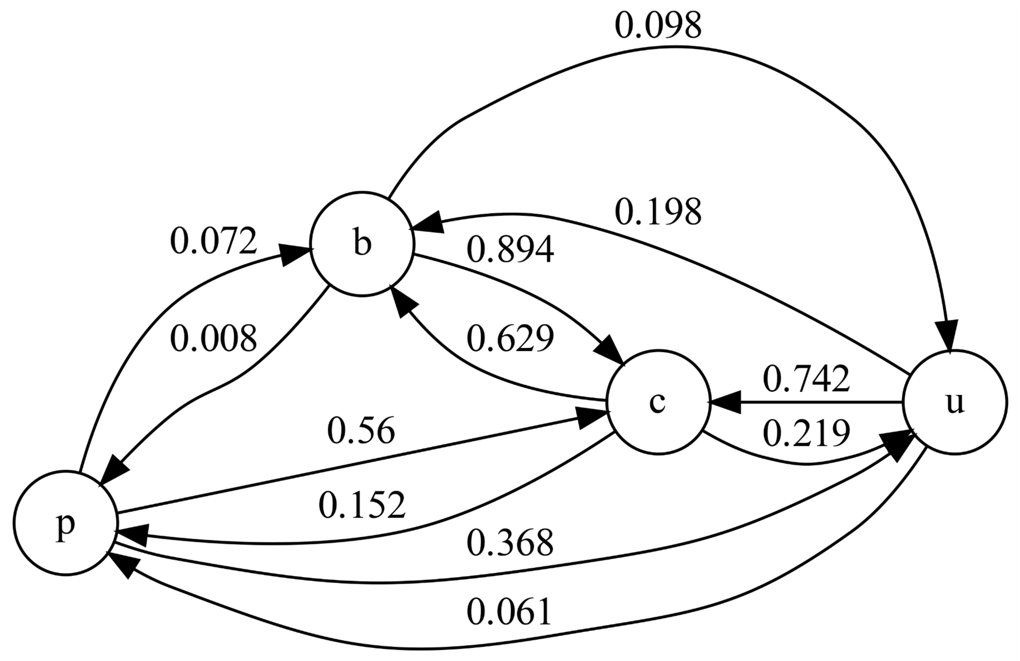


Fig. 3: Markov chain - Transition probabilities of reclassifications

Fig. 1:
- Most reclassified variants have been reclassified once
- Number of variants with at least n reclassifications is exponentially distributed

Fig. 2:
- Reclassifications between ClinVar releases are relatively unlikely

Fig. 3:
- Variants are most likely reclassified into conflicting
- Conflicting variants are most likely reclassified into benign or uncertain

**Result:** ClinVar curators might face a growing set of variants with conflicting interpretations if current reclassification dynamics persist


**Andreas Ruscheinski, Anna Lena Reimler, Roland Ewald, Adelinde M. Uhrmacher**

**Institute for Visual and Analytic Computing | Modeling and Simulation Group**
**University of Rostock, Albert-Einstein-Str. 22, 18059 Rostock, Germany**
